# Content-aware image restoration improves spatiotemporal resolution in luminescence microscopy

**DOI:** 10.1101/2022.04.29.490012

**Authors:** Tobias Boothe, Mario Ivanković, Markus A. Grohme, Jochen C. Rink

## Abstract

Luminescence microscopy is a powerful tool in biomedical imaging applications due to its intrinsically high signal to noise ratio. However, luminescence signal detection requires longer exposure times than fluorescence imaging and is consequently less suited for applications requiring high temporal resolution or throughput. Here we demonstrate that content-aware image restoration can drastically reduce the exposure time requirements in luminescence imaging, thus overcoming one of the major limitations of the technique.

## Introduction

In biomedical sciences, fluorescence microscopy is widely used for the specific visualisation of proteins, organelles, cells, organs or entire organisms^1^. Despite its versatility and applicability on many scales, this imaging technique exposes the sample to potentially harmful excitation light and is also sensitive to artefacts caused by autofluorescence.^1^ In contrast, bioluminescence imaging exploits the light emitted by a chemical reaction between a luciferase enzyme and its substrate (luciferin). Luciferase enzymes originate from approximately 10,000 bioluminescent species across the tree of life and well characterised natural and various biotechnologically optimised or designed luciferases are available as reporters^2^. Luminescence microscopy does not require excitation light and is highly specific due to the practical absence of spontaneous photon emission in biological samples. The technique is therefore especially powerful for imaging photosensitive or highly autofluorescent samples^3,4^. However, a major drawback of bioluminescence imaging are the low signal intensities, which typically require much longer exposure times in comparison with fluorescence imaging. This practically restricts luminescence microscopy applications to immobile samples and imposes throughput limits on high content screening applications. Although shorter exposure times could, in principle, remedy these shortcomings, the inevitable decrease in the signal to noise ratio limits the practical utility of this approach. Recently, Weigert et al. presented an approach for denoising fluorescent microscopy data by utilising deep neural networks, through which signal to noise ratios could be enhanced post acquisition^5^. Here we show that this content-aware image restoration (CARE) can similarly restore luminescence recordings without compromising image quality, allowing exposure time reductions up to 100-fold. By overcoming one of the major limitations of luminescence imaging, our results demonstrate an expanded practicability of luminescence microscopy.

## Results

Convolutional networks have demonstrated strong denoising capabilities in fluorescence microscopy^5,6^. To explore their corresponding utility in luminescence microscopy, we trained a CARE network with luminescence recordings of different exposure times. In our experimental setup, human tissue culture cells expressing untargeted NanoLuc luciferase required 60 s exposure time on a commercially available luminescence imaging system to achieve satisfactory signal to noise ratios. These recordings served as ground truth for restoring signals from exposure times as short as 0.5 s. Training networks on short and long exposure image pairs denoised and restored short exposure images to virtually ground truth quality (Figure 1a, S1a). The restorations from such recordings display a mean absolute error (MAE) that is similar to the technical noise between two subsequently acquired images at 60 s exposure time.

**Figure 1:**
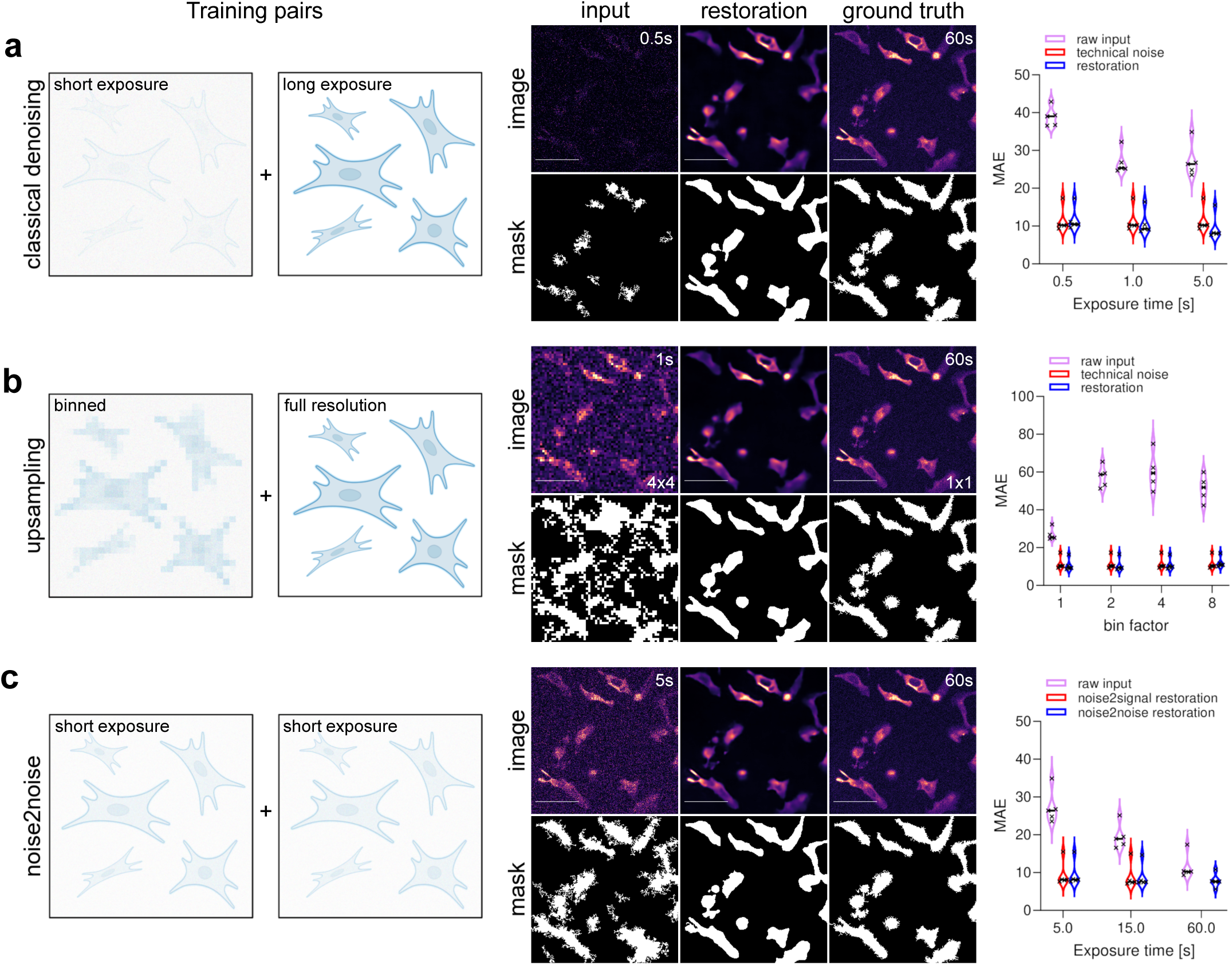
Luminescence image restoration. **(a)** Content-aware image restoration (CARE) allows the restoration of images obtained with short exposure times to images with a signal to noise ratio that is normally achieved with much longer exposure times. **(b)** CARE can be used to upsample and restore binned images to full resolution. **(c)** CARE can be utilised for noise2noise image restorations by training a network with image pairs of short exposure times only. **(a-c)** Schematics for CARE training pairs are illustrated. Restorations were performed on previously unseen data. Binary images represent automatically segmented objects from the respective micrograph. The mean absolute error (MAE) between object masks was used as a quantitative readout to compare restorations with ground truth (lower is better). Scale bars = 10 um

Another commonly used strategy for decreasing exposure times is pixel binning on the camera chip, which sacrifices resolution for signal strength. CARE has previously shown strong capabilities in restoring undersampled z-resolution in 3D recordings.^5^ We sought to transfer this 1D Z-resampling power to the 2D XY-dimension by training a CARE network with binned recordings (2×2, 4×4, 8×8) at low exposure times and unbinned long exposure time ground truth images. We demonstrate that the resolution lost by pixel binning can be restored even from 8×8 binned recordings (Figure 1b, S1b). Therefore, pixel binning and subsequent image restoration provide a further layer of reducing luminescence exposures, with the additional benefit of enhanced visibility of the residual signal in the raw images at short exposures.

Thus far, we described approaches that require long exposure time recording for generating ground truth training data. Technical or biological constraints, however, can make long exposures problematic. Therefore, we additionally explored the capabilities of noise2noise image restoration in which only noisy image pairs are used as training data^7,8^. From such training pairs the CARE network is able to identify and remove statistical noise without the need for long exposure images as ground truth. In our setup exposure times as low as 5 s per image were sufficient to create training data that allowed signal restoration to a degree comparable to that of 60 s exposure times (Figure 1c, S1c). Overall, our data demonstrate the dramatic shortening of luminescence exposure times that are achievable with CARE.

Image restorations performed with CARE - just like any machine learning algorithms - perform best when the training data accurately represents the data to be restored^5^. This often entails frequent retraining of the models whenever experimental conditions change and the recording of training data can thus quickly become a bottle-neck. We therefore evaluated the co-transfection of fluorescently labelled proteins requiring short exposure times as ground truth for their luminescent labelled equivalent. To test this training approach we transiently co-transfected cells with NanoLuc tagged Histone 2B (H2B-Nluc) and eGFP-fused H2B plasmid DNA. We continued to compare the restoration quality of the CARE networks when training with image pairs of short exposure Nluc-H2B signal and eGFP-H2B signal or long exposure Nluc-H2B signal as ground truth respectively. When analysing the by-pixel signal correlation between objects segmented from restorations and ground truth signal, we show that restoration from luminescence signals based on training to fluorescent ground truth is virtually indistinguishable to restorations obtained from training to long exposure luminescence ground truth signals (Figure 2a, S2a). Note that luminescence exposure times as short as fluorescent ground truth exposure times provide sufficient signal from the tagged H2B proteins for an accurate restoration, thus making high-throughput applications of luminescence imaging feasible.

**Figure 2:**
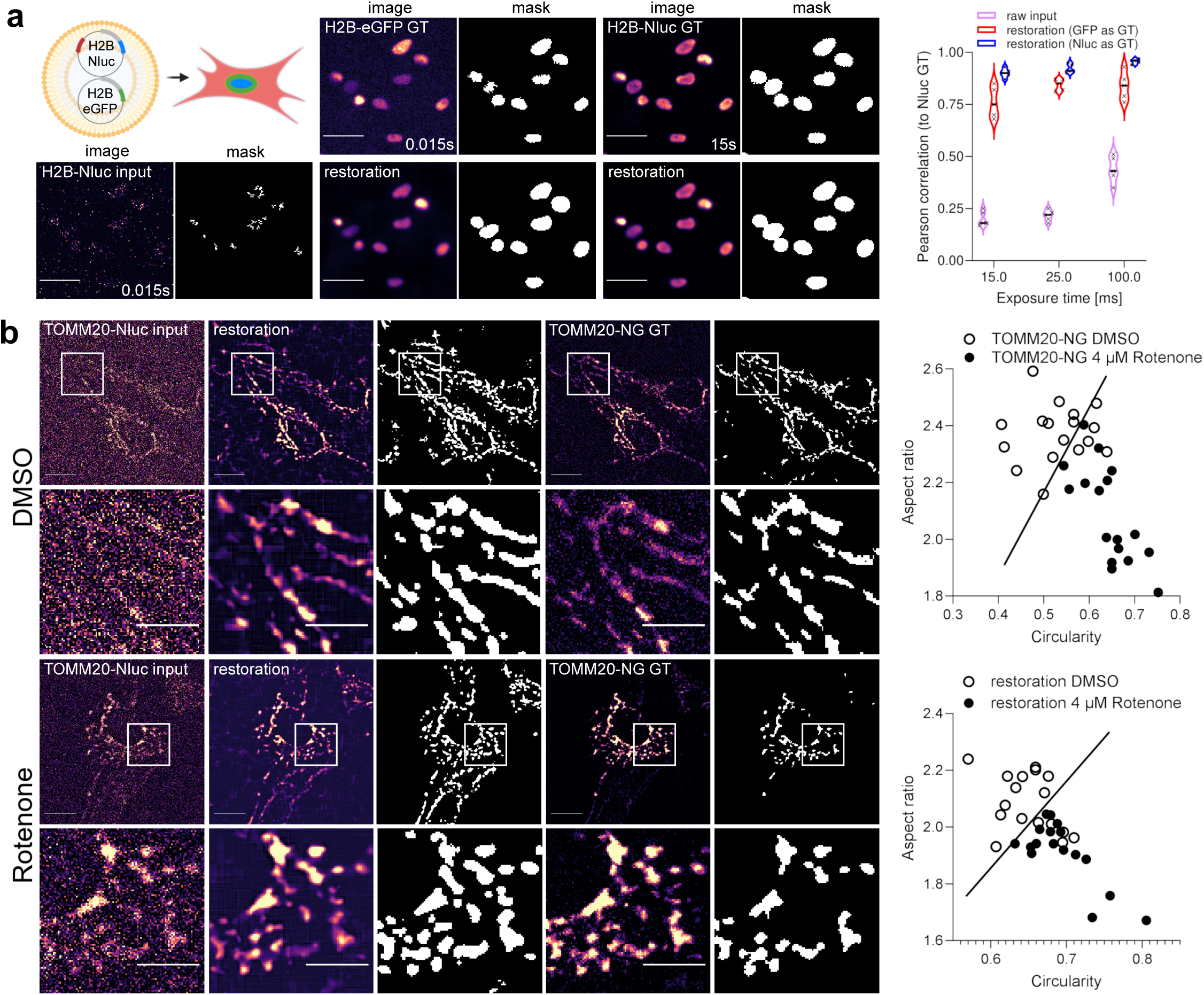
Restoration of luminescence signals to fluorescent ground truth. **(a)** Fluorescently labelled proteins can act as ground truth to reduce exposure times required for training data acquisition. H2B-Nluc and H2B-eGFP were cotransfected and training pairs were generated by capturing short exposure luminescence and short exposure fluorescence signals. Long exposure luminescence signals were recorded for quality control. The Pearson correlation of pixel intensities under masked objects was used as a restoration quality readout. Scale bars = 10 um **(b)** Restoring luminescence signals from diffraction limited structures. Mitochondria were labelled with TOMM20 fusion proteins and a network was trained with luminescence and corresponding fluorescence image pairs. Restorations from luminescence signals allowed for a similar phenotype classification compared to classifications from fluorescently labelled mitochondria. Rotenone treatment was used to induce mitochondrial fragmentation for phenotype scoring. The line in the classification charts delineates all Rotenone treated cells. Scale bars = 5 um

Due to the rather low signal emission, luminescence microscopy has been predominantly applied to whole tissue or organism imaging since the signals are often too weak for the live imaging of subcellular dynamics. To assess restoration qualities in this context, we fused NanoLuc to the outer mitochondrial membrane protein Tomm20^9^ (TOMM20-Nluc) and used mitochondrial dynamics as model system^9^. Due to the continuous remodelling of the mitochondrial network, the recording of long exposure ground truth data is practically impossible. As previously shown (Fig 2 a), fluorescently labelled proteins can act as ground truth while keeping ground truth recordings at short exposure times to avoid motion artefacts. Analogous to the H2B-Nluc/eGFP co-transfection (see above), we trained a network on recordings of cells co-transfected with TOMM20-Nluc and TOMM20-NeonGreen. In this approach we demonstrate that luminescence signals obtained from exposure times as short as 4 s can be reliably restored to the fluorescently labelled TOMM20-NeonGreen ground truth equivalent (Figure 2 b). Mitochondria dynamics, which include fusion and fission events, can be a response to cellular stress and are therefore an important indicator of cellular health^**10**^. To test if the restoration quality is sufficient to characterise mitochondrial phenotypes, we exposed cells to Rotenone - a well characterised respiratory inhibitor that induces mitochondrial fragmentation^11^. We analysed mitochondrial morphology by quantifying the organelle’s circularity and aspect ratio. Despite the challenges associated with segmenting diffraction-limited structures on a widefield microscope, we show that restorations of Rotenone treated cells can be distinguished from their DMSO treated controls. Separation of mitochondrial morphology performed equally well on TOMM20-Nluc signal reconstructions as compared to reconstructions based on fluorescent TOMM20-NeonGreen ground truth of the same cell (Figure 2b). Therefore, our results demonstrate that CARE enables luminescent imaging and analysis of dynamic intracellular compartments that were previously inaccessible due to long exposure time requirements.

## Discussion

The present study demonstrates the utility of CARE in luminescence imaging. Specifically, the previously introduced CARE method enabled the reliable restoration and denoising of luminescence micrographs with initially low contrast, thus shortening required exposure times dramatically. We successfully applied the original CARE network in noise2noise restorations, which is useful when long-term exposures of ground truth are not possible due to biological or technical constraints. Especially in non-dedicated luminescence imaging setups, light pollution by external sources such as instrument LEDs or room light can impact long term exposures and thus greatly complicate the acquisition of the ground truth data.

We further showed that the CARE network architecture can be used to reliably upsample binned images, thus lifting the constraint previously imposed on image resolution. Furthermore, upsampling is useful, when camera read out times are a constraining factor in highly dynamic events as the camera chip read out times for binned images are much shorter than those at full resolution. This is especially interesting for rapid imaging in fluorescence microscopy to which this procedure can be equally applied. It is important to stress that deep learning based restoration methods are - like any restoration method - prone to artefacts,^12^ which can be assessed computationally, as outlined in the original CARE publication^5^.

We finally demonstrate that fluorescence signals can also act as ground truth for luminescence signals in particularly sparse and spatially distinct structures. This further shortens exposure times required for training data acquisition. It is especially useful in very challenging samples that require short exposure times already for training data generation because of biological or practical constraints. With CARE we demonstrate one neural network architecture that can be applied to restore signals in luminescence images. In principle, other networks dedicated to image restoration are potentially equally suitable^6,13^.

With our method, we enable luminescence microscopy imaging at exposure times that shorten acquisitions by a factor of 60 or more. This translates into shortening recording times from minutes to seconds or from hours to minutes. Our method lifts the previously major constraint of long exposure times in luminescence imaging, making this technique now suitable for dynamic and high throughput imaging.

## Supplementary figure legends

**Supplementary Figure S1a: Classical denoising for luminescence image restoration**. Content-aware image restoration (CARE) allows the restoration of images obtained with short exposure times as indicated with a signal to noise ratio that is normally achieved with much longer exposure times. Restorations were performed on previously unseen data. Binary images represent automatically segmented objects from the respective micrograph. Scale bars = 10 um

**Supplementary Figure S1b: Upsampling in luminescence image restoration**. CARE can be used to upsample and restore binned images to full resolution. Restorations were performed on previously unseen data. Binary images represent automatically segmented objects from the respective micrograph. Scale bars = 10 um

**Supplementary Figure S1c: noise2noise denoising in luminescence image restoration**. CARE can be utilised for noise2noise image restorations by training a network with image pairs of short exposure times only. Restorations were performed on previously unseen data. Binary images represent automatically segmented objects from the respective micrograph. Scale bars = 10 um

**Supplementary Figure S2a: Restoration of luminescence signals to fluorescent ground truth**. Fluorescently labelled proteins can act as ground truth to reduce exposure times required for training data acquisition. H2B-Nluc and H2B-eGFP were cotransfected and training pairs were generated by capturing short exposure luminescence and short exposure fluorescence signals. Long exposure luminescence signals were recorded for quality control. Restorations were performed on previously unseen data. Binary images represent automatically segmented objects from the respective micrograph. Scale bars = 10 um

## Online Methods

### Plasmids and Molecular Cloning

DNA constructs containing open reading frames (ORFs) encoding H2B-Nluc and Tomm20-Nluc C-terminal fusion proteins were commercially synthesized (Eurofins Genomics). The ORFs were cloned into pBI-CMV4 (Takara Bio) utilizing 5’-NheI and 3’-SalI restriction sites with reagents from New England Biolabs following standard procedures yielding the final plasmids used for subsequent transfections.

Cytoplasmic Nanoluc was expressed by transfecting U2OS cells with pcDNA3.1-NL plasmid. Plasmid pcDNA3.1-NL was obtained from Addgene (Addgene plasmid #113442). Fluorescently labeled H2B was expressed by transfecting pEGFP-N1-H2B plasmid. H2B-GFP was obtained from Addgene (Addgene plasmid # 11680). Fluorescently labeled Tomm20 was expressed by transfecting pN1-TOMM20-mNG plasmid (Addgene plasmid # 129347).

### Cell Culture

All cell lines were cultured at 37 °C, 90% humidity and 5% CO_2_. U2OS cells were cultured in DMEM medium (Gibco Cat# 31885-023) supplemented with 10% v/v FBS (Anprotec Cat# AC-SM-0033), 100 U/ml Penicillin-Streptomycin (Gibco Cat# 15140-122). Hela CCL-2 cell lines were cultured in DMEM media (Corning Cat# 15-013-CV) supplemented with 10% v/v FBS, 100U/ml Penicillin-Streptomycin and 10 mM L-glutamine (Gibco Cat# 25030-024).

### Transfections

All plasmid transfections were performed at 70% confluency using Lipofectamine 3000 transfection reagent using 2 µl Lipofectamine and 2 µl P3000 reagent/µg DNA (Thermo Fisher Scientific). For transfecting 30 mm dishes or 75 cm^2^ flask, a total of 2.5 µg or 25 µg of plasmid DNA were used, respectively. For double transfections, plasmids were combined equally while retaining absolute amounts used for single transfections.

### Fluorescence-activated cell sorting

To ensure coexpression in experiments that were used to train luminescence signal on fluorescent ground truth signal, luminescence fusion proteins were cloned into a bidirectional vector (pBI-CMV4) that also expresses dsRed2 as an expression control for the luminescent fusion protein (see section ‘plasmids and cloning’ for details).

For experiments in which co-expression of fluorescently labeled H2B and Nluc conjugated H2B was required, a transfected, confluent 75 cm^2^ flask of HelaCCL cells was harvested. Cells were resuspended in complete DMEM and kept on ice and FACsorted at 5 °C: eGFP^+^/dsRed2^+^ double positive populations were isolated by FACS using a SONY Cell Sorter SH800 with a 100µM microfluidics sorting chips. Scatter characteristics utilizing FSC-A/BSC-A were used to exclude debris and FSC-A/FSC-W to exclude doublets. eGFP^+^/dsRed2^+^ cells were sorted using the “Semi-Purity” sort mode with a sensor gain of 32% for dsRed2 (FL3-600/60) and eGFP (FL2-525/50). During the sort, 488 nm and 561nm lasers were active. Per 35mm glass bottom dish 300.000 double positive cells with comparatively high expression ratios of both eGFP and dsRed were seeded.

### Luminescence assays

For cells expressing cytosolic NanoLuc or H2B-Nanoluc fusion proteins, Nano-Glo Endurazine (Promega) was used as a substrate at a 1x final concentration. Cells were imaged 1h after addition of the substrate.

For cells expressing Tomm20-Nanoluc fusion protein, Nano-Glo Vivazine (Promega) was used as a substrate at a 1x final concentration. Cells were imaged 1.5 h after addition of the substrate.

### Microscopy

All images were recorded using Olympus’ LV200 bioluminescence imaging platform. eGFP and NeonGreen were excited through a 470/11nm bandpass filter. Emission for eGFP and NeonGreen was collected with a 525/25nm bandpass filter. Luminescence was detected without any emission filter. Olympus’ 20x NA 0.8 UPLXAPO objective was used to image cytoplasmic NanoLuc and H2B-Nluc/eGFP signals. Olympus’ 100x NA 1.5 UPLAPO OHR Objective was used to image Tomm20-Nluc/mNeonGreen. For signal detection, an Andor iXon 888 Ultra EM-CCD camera, deep cooled to -85°C at a 1 Mhz readout rate with an EM gain of 300 was used. Cells were incubated with a Tokai Hit stage top incubator providing full environmental control.

Cells for imaging experiments were cultured in 30mm glass bottom dishes (ibidi, Cat# 81158). Cells were imaged in CO_2_-independent, phenol-red free, L15 Leibovitz media (Thermo Fisher Cat# 21083027) supplemented with 10% v/v FBS and 100U/ml Penicillin-Streptomycin. L15 Leibovitz imaging media for Hela CCL cells was additionally supplemented with 10mM L-Glutamine.

### Data recording, Deep Neural Network training and image restoration

For training, the previously published CSBDeep package v0.6.0 was used (http://csbdeep.bioimagecomputing.com). A detailed documentation of this software is available at http://csbdeep.bioimagecomputing.com/doc. We performed all training and prediction pipelines using our publicly available docker container that can be obtained at https://hub.docker.com/r/tboo/csbdeep_gpu_docker. Generalised Python scripts that can be used with this container for preparing training data, network training and prediction are available at https://gitlab.gwdg.de/rinklab_public/lumicare.

Networks were trained on a Lenovo ThinkSystem SR670 server equipped with two Intel Xeon Gold 6234 CPUs, 768GB RAM and four NVIDIA Tesla V100 32GB GPUs.

Networks were trained for each low-high signal condition separately, providing best restoration performance.

All training data were recorded in 3D (XYZ) and all networks were trained as 3D networks. For acquisition of training data, input (low signal) condition(s) and ground truth/target condition (high signal) were imaged consecutively per plane before proceeding to the next z-plane. Training data for noise-to-noise training was obtained by taking 2 consecutive images with identical imaging parameters. To train networks for upsampling, training data was obtained by software binning using the “bin” function in Fiji. These binned images were subsequently upsampled without interpolation to match the pixel dimensions of the respective ground truth image. To increase training data complexity, each raw stack was rotated 3 times in 90° increments and subsequently each of the resulting stacks was mirrored horizontally resulting in an 8-fold increase of available training data. The table below summarises the key parameters used for training data acquisition, training data preparation and network training.

|  |  | Structures to restore |  |  |
| --- | --- | --- | --- | --- |
|  |  | Cytoplasmic Nluc | H2B-Nluc | Tomm20-Nluc |
| Microscope parameters for generating raw training data | XYZ camera ROI (pixels, unbinned) | 1024x1024x16 | 512x512x16 | 512x512x16 |
|  | Z stepping | 570 nm | 570 nm | 130 nm |
|  | Number of Stacks | 8 | 14 | 90 |
|  | Objective | 20x XAPO | 20x XAPO | 100x APO |
| Processed Training data | Patch dimensions XYZ (pixels) (patch_size) | 64x64x16 | 64x64x16 | 64x64x16 |
|  | number of randomly sampled patches per raw image stack (n_patches_per_image) | 200 | 250 | 150 |
| Training parameters | Number of epochs (train_epochs) | 150 | 100 | 200 |
|  | Steps per epoch (train_steps_per_epoch) | 200 | 400 | 600 |
|  | Batch size (train_batch_size) | 128 | 128 | 128 |
|  | Fraction used for validation (validation_split) | 0.1 | 0.2 | 0.3 |

Networks were trained with probabilistic per pixel prediction (`probabilistic`=`True`). All other network parameters were used in default settings.

For all conditions, the trained models were used to restore images by tiling the respective stacks 2×2×2 in XYZ using the csbdeep API.

### Image analysis

All image processing and analysis was performed using the ImageJ distribution Fiji v2.3.0 ^14^. To quantify the disagreement between ground truth images of cells expressing cytoplasmic Nanoluc and respective restoration results from low signal images, the mean absolute error (MAE) between cell masks was determined using the SNR plugin v06.05.2011^15^. To obtain the masks, all images were converted to 16bit, thresholding was applied using the “Mean” auto thresholding function with enabling the “dark background” option. Resulting particles were filtered by size (>100 pixel). The resulting masks were inverted and used with the SNR plugin to quantify the MAE between the ground truth masks as the reference images and the corresponding restoration or input masks as test images.

To quantify the disagreement between ground truth images of cells expressing labeled H2B and respective restoration results from low signal images, the Pearson correlation between intensities of masked nuclei was determined. For that purpose, ground truth images were thresholded using the “Triangle” auto thresholding function enabling the “dark background” option. Resulting particles were filtered by size (>5 pixel). The resulting masks were inverted and used as masks for determining the Pearson correlation of raw pixel values between an input/restoration image and the respective ground truth images utilizing the “Coloc2” plugin v3.0.5 in Fiji.

For quantifying mitochondrial morphology all images (including restorations) were maximum projected along Z and a rolling ball background subtraction (radius = 5 pixel) was applied using Fiji. To quantify the disagreement between ground truth images of cells expressing labeled Tomm20 and respective restoration results from low signal images, a pixel classifier to segment mitochondria was trained using ilastik software v1.3.3 ^16^. For training this pixel classifier, a subset of mitochondria and background was annotated in 10 images total (5 DMSO control, 5 Rotenone treated) of cells expressing Tomm20-NeonGreen (ground truth images). This trained classifier predicted accurate mitochondria masks in all images used for analysis. The resulting masks were subsequently analyzed for their morphology with Fiji by setting an object size threshold (>20 pixel). Circularity and aspect ratio of these segments was measured via the ‘shape’ measurements module. The values displayed in Figure 2 are object measurement mean values per image analysed.

## Acknowledgements

The authors would like to thank Britta Schroth-Diez and Jan Peychl from the Light Microscopy Facility at the Max Planck Institute for Molecular Cell Biology and Genetics for generous support and encouragement during the first proof of concept experiments for this study. We would further like to thank Dajana Burghardt for technical assistance with molecular cloning.

pcDNA3.1 NL was a gift from Erich Wanker. pEGFP-N1-H2B was a gift from Geoff Wahl. pN1-TOMM20-mNG was a gift from Yasushi Okada. Mammalian cell lines were a gift from Stefan Jakobs.

