## Supplementary figures and images for "Content-aware image restoration improves spatiotemporal resolution in luminescence microscopy"

### Supplementary Figure S1a

#### Supplementary Figure S1a

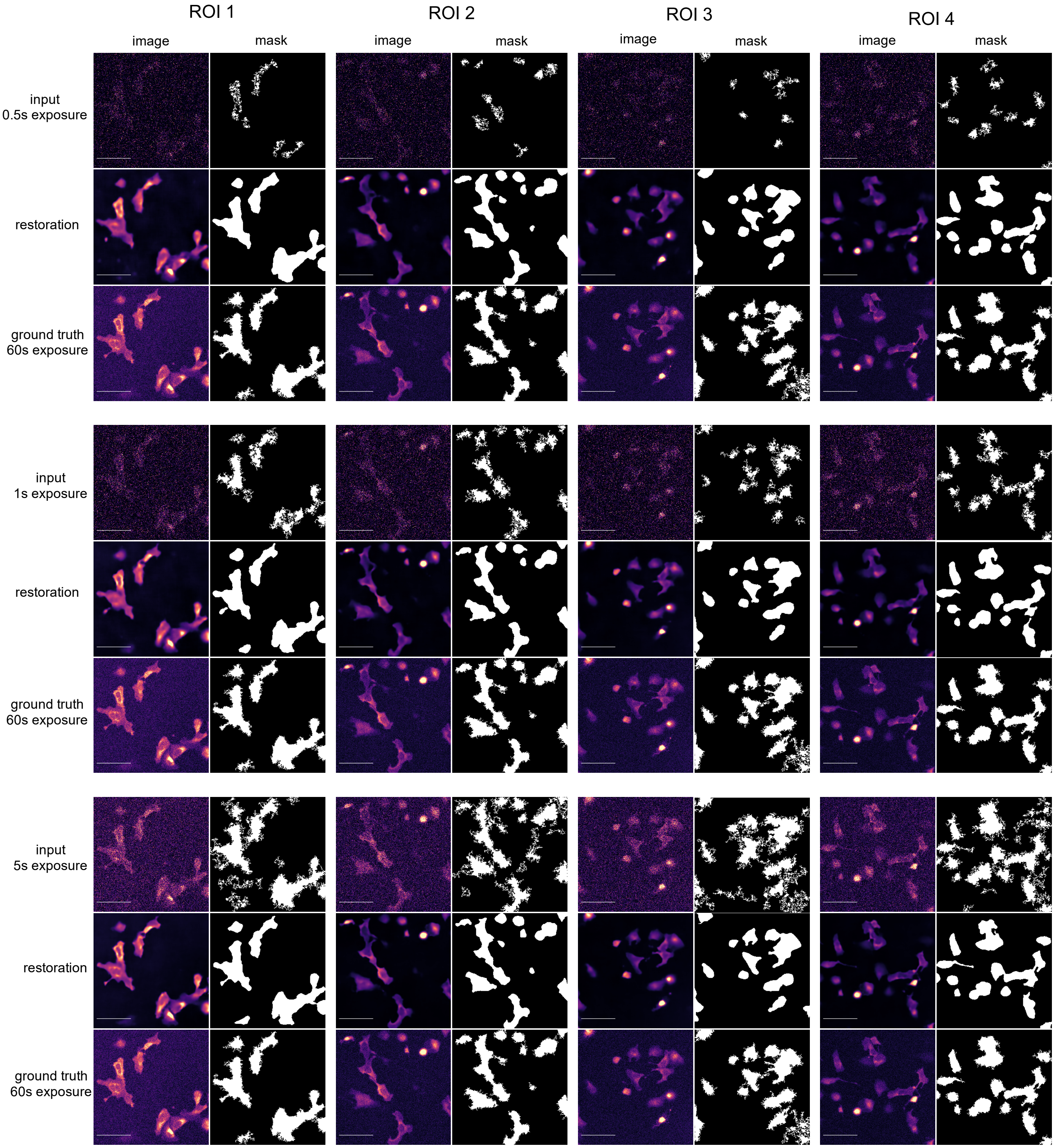

### Supplementary Figure S1b

#### Supplementary Figure S1b

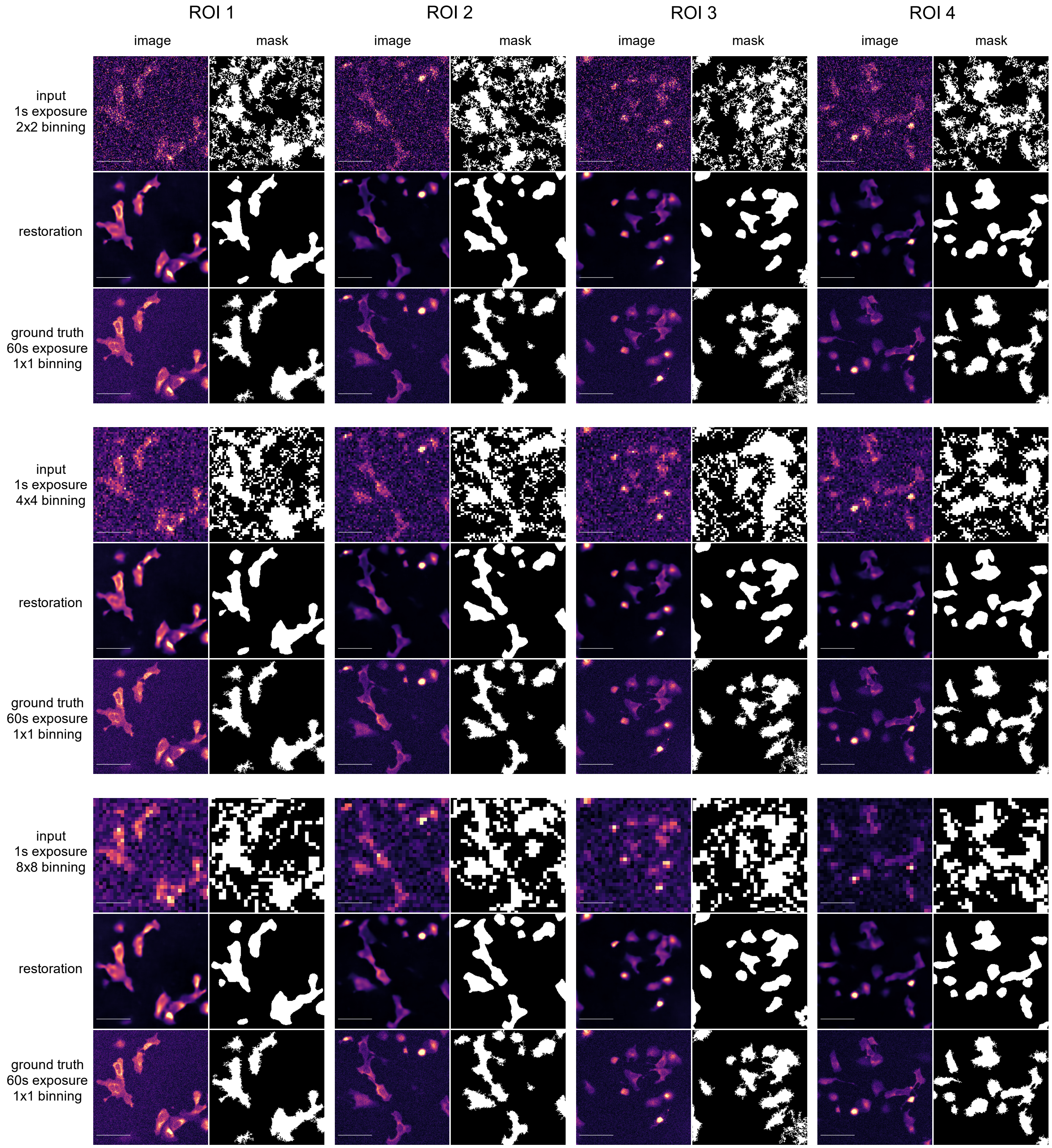

### Supplementary Figure S1c

#### Supplementary Figure S1c

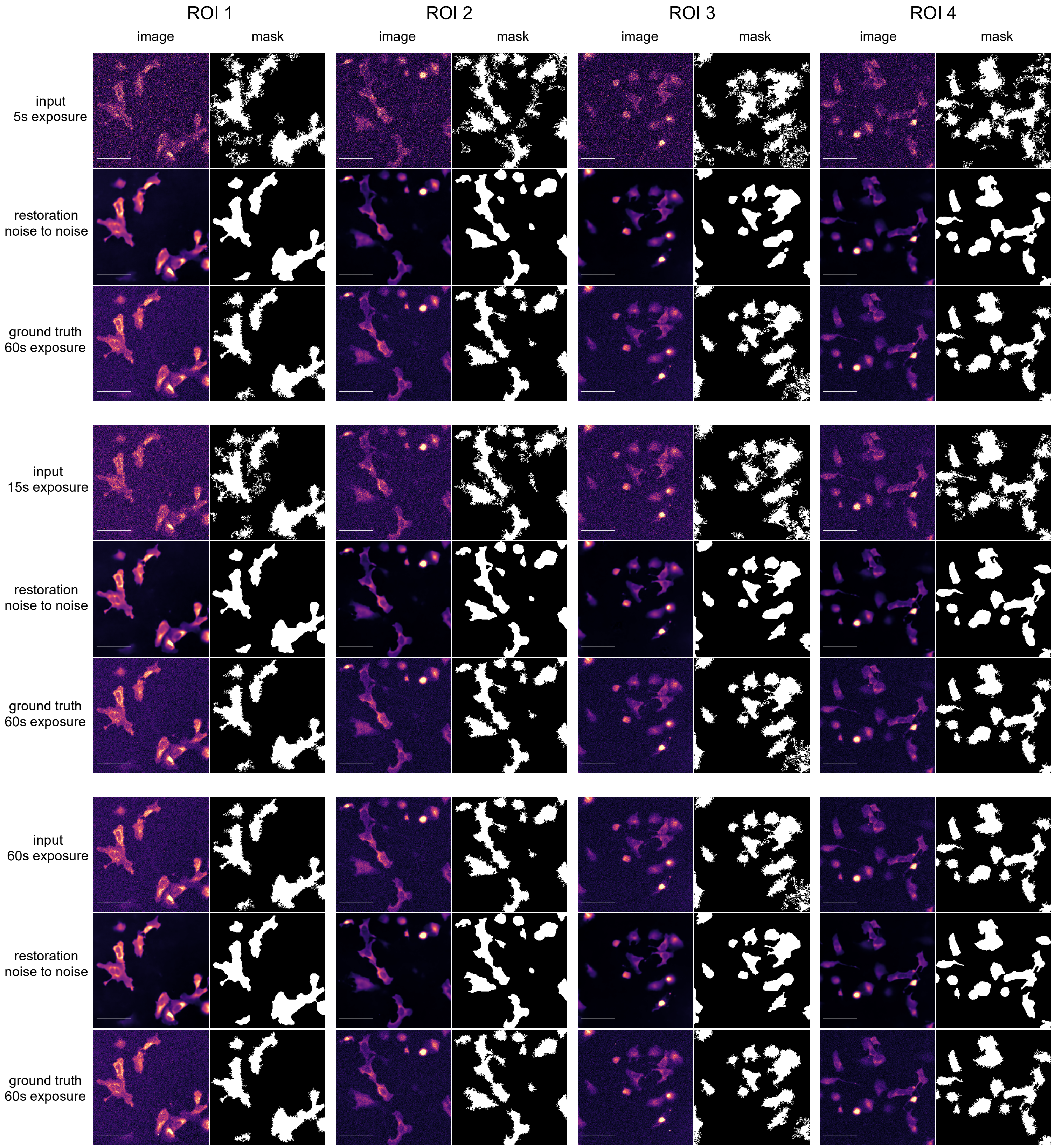

### Supplementary Figure S2a

#### Supplementary Figure S2a

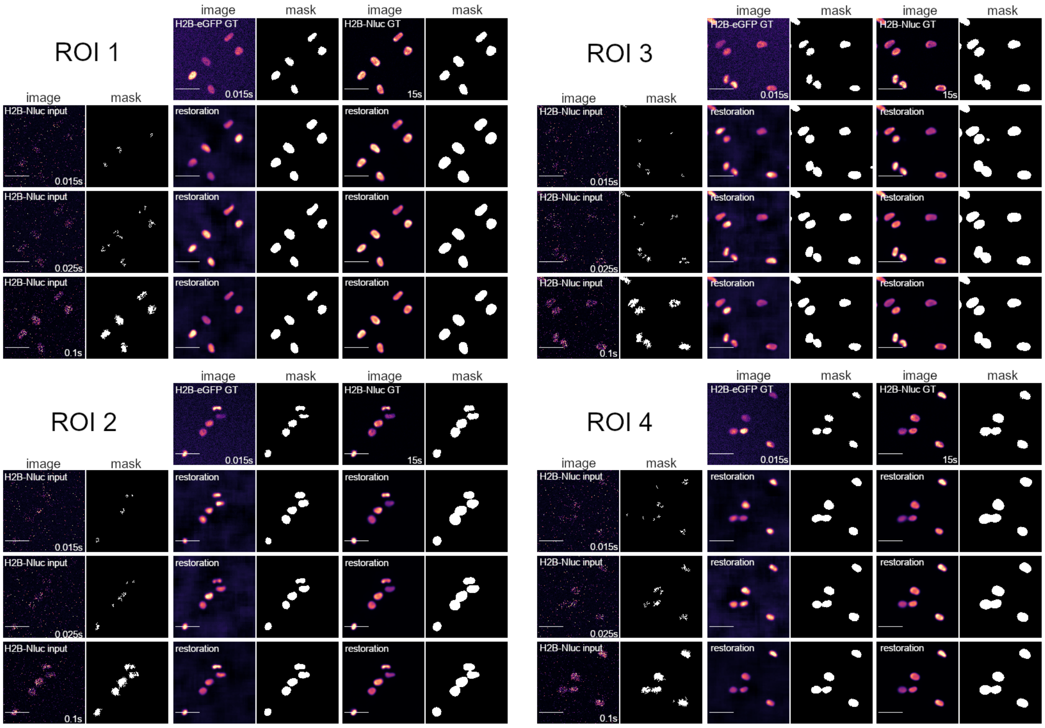
